## Supplemental Information for "Watching the release of a photopharmacological drug from tubulin using time-resolved serial crystallography"

**This PDF file includes:**

Supplementary Figures. S1 to S6
Supplementary Tables S1 to S2

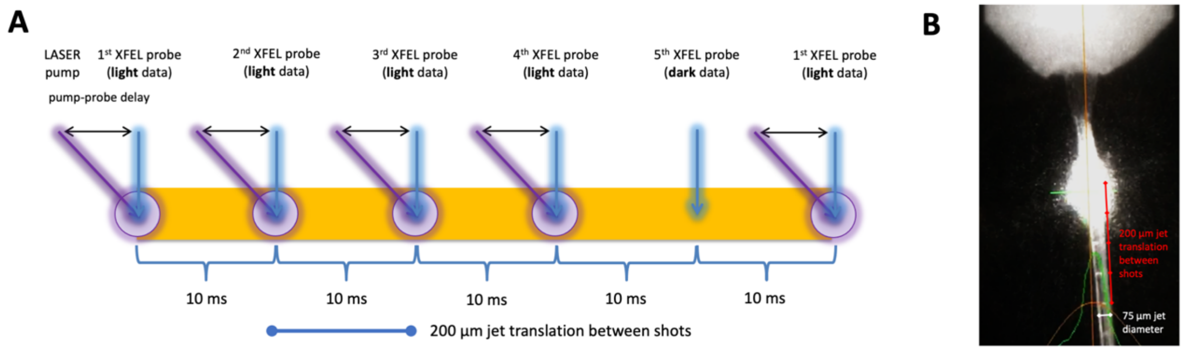


**Supplementary Figure 1: Experimental setup.** (**A**) The 4:1 pump-probe scheme used with the high-viscosity sample extruder[45] we introduced for crystal delivery time-resolved crystallography experiments[46] is depicted. Distance between shots taken at 100 Hz was about 200 µm and the laser spot size ($1/e^2$) was about 65 µm (depicted as purple circle). Every $5^{th}$ pattern was taken without the laser turned on. In this setup each time delay was collected in about 50 minutes measuring time and needed approximately 5 mg of tubulin. Notably, we suggest to optimize sample injection before the experiment to increase sample and time efficiency during the beamtime[47]. Alternative sample delivery methods like solid-supports[48] or tape-drives[49] could be used to target longer time delays and thus measure ligands with longer residence times. (**B**) Image showing the laser on the jet and the penetration marks of the XFEL. Taking the 75 µm jet thickness to provide a scale in the picture it is evident that individual shots which are visible as white lines on the jet are evenly spaced. The laser glow in the image is overloaded and cannot be used to draw conclusions on the focal spot size of the laser, which was accurately measured using a knife edge scan at the sample position.

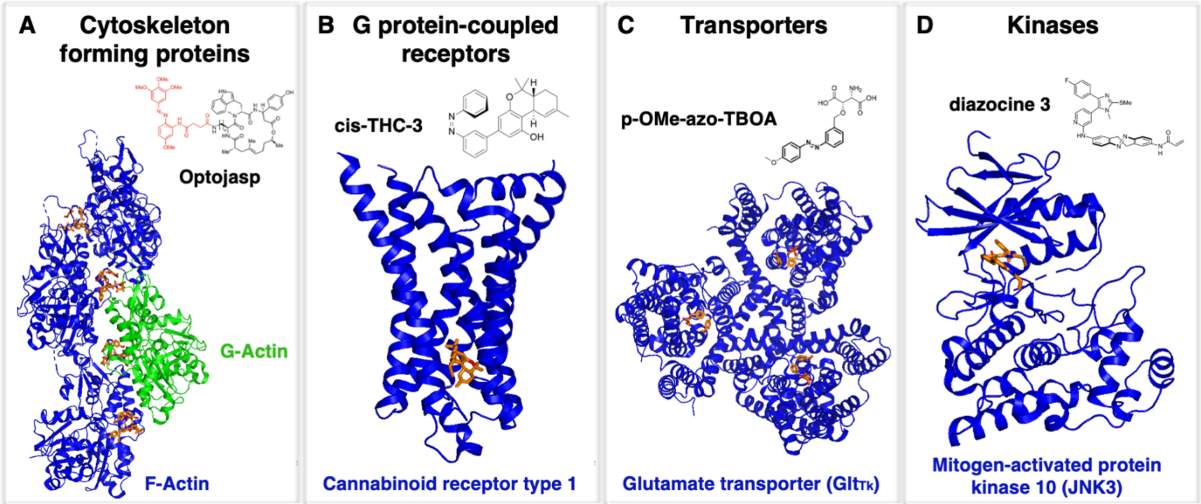


**Supplementary Figure 2: Selection of protein classes that have been targeted with photopharmacological compounds.** (**A**) A class of cytoskeletal proteins: Actin. Most abundant cellular protein and an essential component of the eukaryotic cytoskeleton[50, 51, 52]. (**B**) A class of G protein-coupled receptors: Cannabinoid receptor type 1. One of the most widely expressed GPCR in the central nervous system. Its activation is associated with mood, motor coordination, memory, and recognition[53, 54]. (**C**) A class of membrane transporters: Glutamate transporter $Glt_{Tk}$. Disruption linked to neurotoxicity under ischemic conditions and epilepsy[55]. (**D**) A class of signaling kinases: Mitogen-activated protein kinase 10 (JNK3). A key signaling enzyme in the cellular stress response, targeted for the treatment of neurodegenerative diseases including Alzheimer's, Huntington's, and Parkinson's disease[56]. These examples suggest that the approach to employ photochemical affinity switches to study protein-ligand interaction dynamics could be expanded to a series of important targets from pharmacologically relevant protein families.

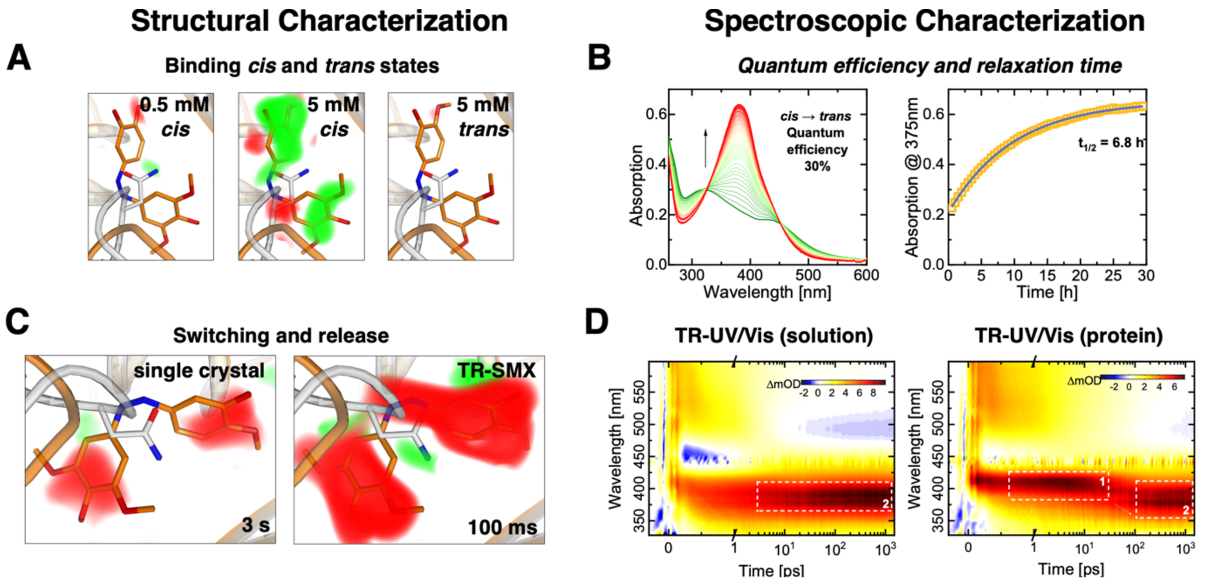


**Supplementary Figure 3: Initial characterization of the azo-CA4 photoswitch. (A)** Soaking of tubulin crystals with 0.5 mM *cis*-azo-CA4, 5 mM *cis*-azo-CA4, or 5 mM *trans*-azo-CA4. The binding pocket remained unoccupied up to azo-CA4 concentrations of 5 mM when the ligand was not exposed to light, suggesting that a light-triggered release of azo-CA4 should be feasible. (**B**) Titration experiments used to determine the quantum efficiency of conversion and relaxation of azo-CA4 over time in crystallization buffer. The quantum efficiency of 30% is large enough to reach sufficient activation levels in time-resolved serial crystallography experiments. We further confirmed that the *cis*-to-*trans* thermal relaxation is slow enough to maintain the *cis* conformation without constant illumination during a single sample injection of about 30 minutes. (**C**) Ligand release after photoinduced isomerization followed by a brief annealing of a single cryo-cooled crystal soaked with 1.25 mM of *cis*-azo-CA4, and ligand release after 100 ms in a time-resolved serial crystallography experiment. A single tubulin crystal was soaked with 1.25 mM of *cis*-azoCA4 before collecting a conventional cryo-dataset. Light data was collected by blocking the cryo-stream and illuminating the crystal before collecting a second dataset on a crystal position ~150 µm away from the first position. The isomorphous difference map between both datasets showed that azo-CA4 is released from the crystal upon photoswitching. We then moved on to a time-resolved serial crystallography experiment at the synchrotron and found that the ligand was fully released from its tubulin-binding site after 100 ms of illumination. (**D**) Time-resolved UV-Vis spectroscopy revealed that tubulin modifies the compound relaxation behavior up to the late picosecond range, suggesting that the ligand isomerization inside the binding pocket is finished after 1 ns. All panels show isomorphous difference maps that are displayed in red (negative) and green (positive) at 3 σ. The substrate free apo form of tubulin is shown in grey and the substrate bound form in orange.

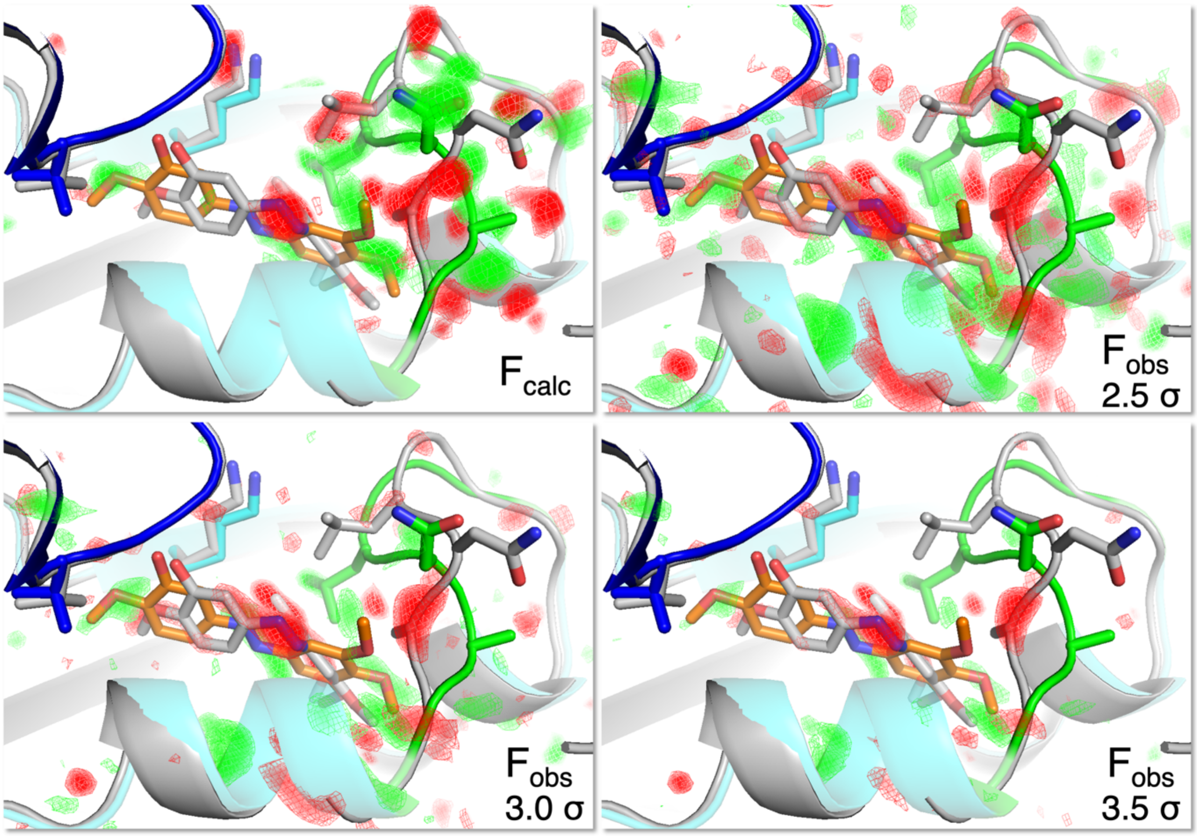


**Supplementary Figure 4: Comparison of $F_{obs}$ and $F_{calc}$ difference maps.** Experimentally observed $F_{obs}$(1ms)-$F_{obs}$(dark) and calculated $F_{calc}$(1ms-model)-$F_{calc}$(dark-model) difference density maps within the ligand binding site are shown. Positive and negative difference densities are displayed in green and red, respectively. The initial dark model is shown in grey and the 1 ms structure as blue (α tubulin), cyan (β tubulin), and green (βT7 loop). Selected side chains are displayed in stick representation. The $F_{calc}$ difference density map (top) was calculated using B-factors of 30 for all atoms and is displayed at a sigma level of 3.5 sigma, while the $F_{obs}$ maps are shown at levels from 2.5 to 3.5 sigma. The $F_{calc}$ maps represent a full activation level and data without noise and even though this prevents direct comparison of sigma levels, there is an excellent agreement between the simulated and calculated difference maps.

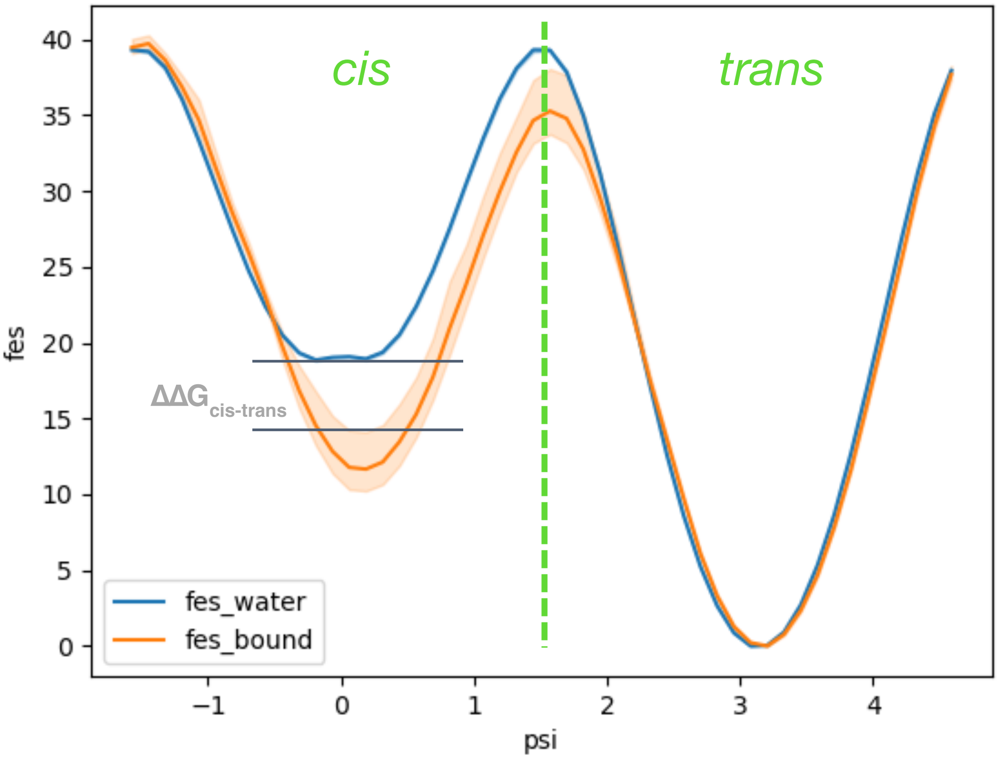


**Supplementary Figure 5: Computational analysis of the binding energy landscape provides a rationale for how cis-trans isomerization initiates ligand unbinding.** The plot indicates the energy gain of the *cis* vs. the *trans* azo-CA4 isomer upon binding to tubulin. A single WTM simulation was carried out with a custom protocol to sample conformations compatible to the XFEL-based time-resolved structures. Block analysis with block size of 50 ns was performed to assess convergence and to plot 95% confidence interval error bands around converged values.

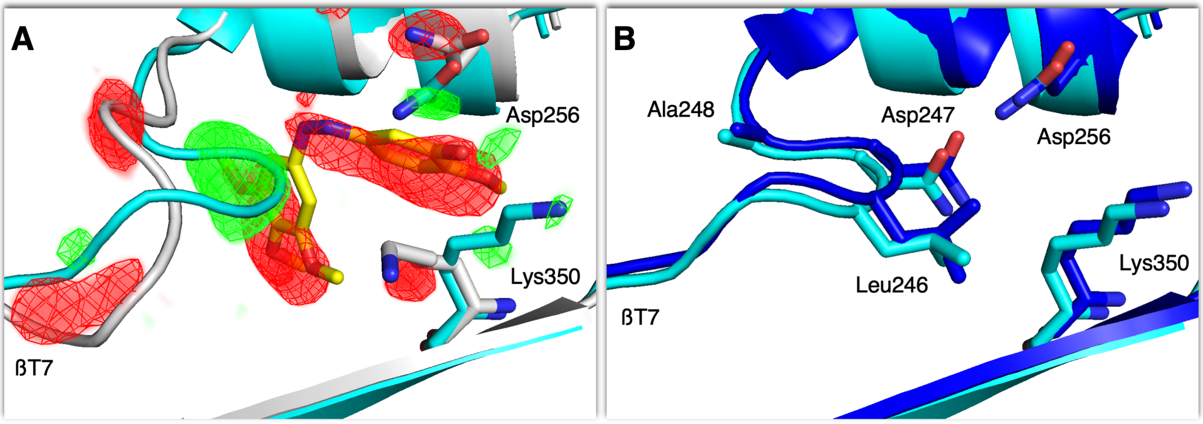


**Supplementary Figure 6: Comparison of tubulin-azo-CA4 complex after photoinduced ligand release with a structure of apo-tubulin**. (**A**) Overlay of liganded dark (grey) and 100 ms (cyan) tubulin structures obtained by serial synchrotron crystallography. The corresponding difference densities (positive in green, negative in red, $F_{obs}$(100 ms)-$F_{obs}$(dark), sigma level = 5.0) confirm the release of azo-CA4 and reorganization of the colchicine-site in tubulin. (**B**) Overlay of the 100 ms structure (cyan) and the one determined using data from apo-tubulin crystals (blue).

### Supplementary Table 1: Crystallographic data statistics

| | **Dark (XFEL)** | **1 ns (XFEL)** | **10 ns (XFEL)** | **100 ns (XFEL)** | **1 ms (XFEL)** | **10 ms (XFEL)** | **100 ms (XFEL)** | **1 ms (XFEL)** | **10 ms (XFEL)** | **Apo (XFEL)** | **100 ms (SYN)** | **Dark (SYN)** |
|---|---|---|---|---|---|---|---|---|---|---|---|---|
| **Data collection** | | | | | | | | | | | | |
| **Space group** | $P2_1$ | | | | | | | | | | | |
| *a*, *b*, *c* (Å) | 74.5, 92.6, 84.0 | | | | | | | | | 74.7, 92.7, 84.1 | 74.3, 91.9, 83.7 | |
| a, b, g (°) | 90, 96.7, 90 | | | | | | | | | 90, 96.4, 90 | 90, 96.82, 90 | |
| **Indexed patterns** | 370356 | 63970 | 69058 | 61091 | 73798 | 67496 | 57915 | 60644 | 89399 | 82666 | 41314 | 102238 |
| **Indexing rate (%)** | 16.6 | 23.3 | 18.1 | 19.8 | 13.2 | 23.5 | 31.2 | 16.7 | 22.6 | 25.6 | 18.1 | 16.5 |
| **Overall statistics: 11.09 Å – 1.70 Å (high-resolution statistics: 1.76 Å – 1.70 Å)** | | | | | | | | | | **11.09-1.80 Å (1.86-1.80 Å)** | **92.11-2.1 Å (2.18-2.10 Å)** | |
| **No. reflections** | 123960 (12356) | 122155(12154) | 122150 (12154) | 122150 (12154) | 122150 (12154) | 122154 (12154) | 122150 (12154) | 122153 (12154) | 122152 (12154) | 104994 (10494) | 38945 (117) | 43965 (945) |
| **Completeness (%)** | 100 (100) | 100 (100) | 100 (100) | 100 (100) | 100 (100) | 100 (100) | 100 (100) | 100 (100) | 100 (100) | 100 (100) | 60 (2) | 67 (15) |
| **Multiplicity** | 2089.8 (1000.6) | 365.9 (188.4) | 347.5 (175.9) | 314.1 (164.8) | 404.2 (208.4) | 392.6 (207.5) | 309.1 (155.8) | 328.9 (161.2) | 586.1 (313.1) | 429.5 (194.3) | 274.2 (216.6) | 602.5 (504.6) |
| **$R_{split}$ (%)** | 7.5 (78.85) | 16.9 (217.8) | 17.2 (204.3) | 18.1 (180.4) | 16.6 (268.7) | 16.7 (174.4) | 17.5 (299.0) | 18.1 (253.1) | 13.8 (106.5) | 14.2 (372.3) | 10.5 (69.48) | 8.0 (49.9) |
| **$CC_{1/2}$** | 0.992 (0.694) | 0.971 (0.265) | 0.968 (0.274) | 0.964 (0.323) | 0.972 (0.210) | 0.971 (0.343) | 0.969 (0.170) | 0.968 (0.217) | 0.979 (0.558) | 0.981 (0.14) | 0.992 (0.528) | 0.996 (0.660) |
| **$\langle I/\sigma(I)\rangle$** | 9.53 (1.43) | 4.01 (0.50) | 4.16 (0.56) | 4.05 (0.62) | 4.05 (0.43) | 4.26 (0.64) | 3.68 (0.38) | 3.79 (0.45) | 5.36 (1.03) | 4.59 (0.32) | 7.87 (1.25) | 10.09 (1.62) |
| **PDB Code** | 7YYQ | 7YYV | 7YYW | 7YYX | 7YYY | 7YYZ | 7YZ0 | 7YZ1 | 7YZ2 | 7YZ3 | 7YZ5 | 7YZ6 |

### Supplementary Table 2: Refinement statistics

| | Dark (XFEL) | 1 ns (XFEL) | 10 ns (XFEL) | 100 ns (XFEL) | 1 µs (XFEL) | 10 µs (XFEL) | 100 µs (XFEL) | 1 ms (XFEL) | 10 ms (XFEL) | Apo (XFEL) | ~100 ms (SYN) | Dark (SYN) |
|---|---|---|---|---|---|---|---|---|---|---|---|---|
| **Resolution (Å)** | 9.5 Å – 1.7 Å | 9.5 Å – 2.2 Å | 9.5 Å – 2.2 Å | 9.5 Å – 2.2 Å | 9.5 Å – 2.2 Å | 9.5 Å – 2.2 Å | 9.5 Å – 2.2 Å | 9.5 Å – 2.2 Å | 9.5 Å – 2.2 Å | 9.5 Å – 1.80 Å | 73.75 Å – 2.1 Å | 73.75 Å – 2.1 Å |
| **No. reflections** | 123591 (8816) | 53409 (3901) | 53221 (3910) | 52461 (3763) | 52234 (3710) | 53180 (3840) | 52580 (3778) | 53111 (3915) | 53134 (3657) | 102745 (1827) | 38921 (68) | 43943 (652) |
| **$R_{work}$ / $R_{free}$ in %** | 12.13 / 15.38 | 31.11 / 35.19 | 31.13 / 34.33 | 31.10 / 35.53 | 30.46 / 34.88 | 29.87 / 35.48 | 30.48 / 36.29 | 30.18 / 35.17 | 28.52 / 33.36 | 18.45 / 21.57 | 17.28 / 23.08 | 17.76 / 22.64 |
| **No. atoms** | 8482 | 8284 | 8352 | 8211 | 8217 | 8228 | 8262 | 8274 | 8210 | 8358 | 8198 | 8646 |
| **Protein** | 8392 | 7915 | 7945 | 7915 | 7909 | 7907 | 7912 | 7935 | 7931 | 7996 | 7986 | 8392 |
| **Ligands** | 90 | 57 | 57 | 57 | 57 | 57 | 57 | 57 | 33 | 34 | 34 | 90 |
| **Water** | 544 | 284 | 322 | 211 | 223 | 236 | 265 | 254 | 218 | 300 | 150 | 164 |
| ***B*-factors** | 44.82 | 34.34 | 33.25 | 34.64 | 40.97 | 35.09 | 35.52 | 30.69 | 41.84 | 48.25 | 39.94 | 42.16 |
| **Protein** | 44.27 | 34.43 | 33.34 | 34.69 | 41.10 | 35.17 | 35.60 | 30.77 | 41.89 | 48.32 | 39.08 | 42.30 |
| **Ligands** | 36.79 | 26.95 | 32.38 | 29.40 | 32.34 | 28.44 | 31.26 | 24.09 | 28.41 | 32.19 | 21.95 | 36.13 |
| **Water** | 54.51 | 33.47 | 27.06 | 34.37 | 38.62 | 33.88 | 34.30 | 29.50 | 42.22 | 48.88 | 35.76 | 38.15 |
| **Bond lengths (Å)** | 0.008 | 0.001 | 0.001 | 0.002 | 0.002 | 0.001 | 0.001 | 0.004 | 0.001 | 0.003 | 0.001 | 0.002 |
| **Bond angles (°)** | 0.885 | 0.414 | 0.424 | 0.455 | 0.436 | 0.409 | 0.426 | 0.673 | 0.422 | 0.670 | 0.417 | 0.469 |
| **Ramachandran favored / allowed / outliers in %** | 98.71 / 1.29 / 0.00 | 96.73 / 3.17 / 0.10 | 97.23 / 2.57 / 0.20 | 96.93 / 2.77 / 0.30 | 96.73 / 2.87 / 0.40 | 97.23 / 2.57 / 0.20 | 96.82 / 2.68 / 0.50 | 96.04 / 3.47 0.50 | 96.13 / 3.57 0.30 | 98.13 / 1.77 / 0.10 | 95.97 / 3.54 / 0.49 | 97.72 / 2.08 / 0.20 |
| **PDB Code** | 7YYQ | 7YYV | 7YYW | 7YYX | 7YYY | 7YYZ | 7YZ0 | 7YZ1 | 7YZ2 | 7YZ3 | 7YZ5 | 7YZ6 |

77. D.A. Case HMA, K. Belfon, I.Y. Ben-Shalom, S.R. Brozell, D.S. Cerutti, T.E. Cheatham, III, G.A. Cisneros, V.W.D. Cruzeiro, T.A. Darden, R.E. Duke, G. Giambasu, M.K. Gilson, H. Gohlke, A.W. Goetz, R. Harris, S. Izadi, S.A. Izmailov, C. Jin, K. Kasavajhala, M.C. Kaymak, E. King, A. Kovalenko, T. Kurtzman, T.S. Lee, S. LeGrand, P. Li, C. Lin, J. Liu, T. Luchko, R. Luo, M. Machado, V. Man, M. Manathunga, K.M. Merz, Y. Miao, O. Mikhailovskii, G. Monard, H. Nguyen, K.A.

O'Hearn, A. Onufriev, F. Pan, S. Pantano, R. Qi, A. Rahnamoun, D.R. Roe, A. Roitberg, C. Sagui, S. Schott-Verdugo, J. Shen, C.L. Simmerling, N.R. Skrynnikov, J. Smith, J. Swails, R.C. Walker, J. Wang, H. Wei, R.M. Wolf, X. Wu, Y. Xue, D.M. York, S. Zhao, and P.A. Kollman. Amber 21. In: *University of California, San Francisco*) (2021).
